# Physical activity promotes longitudinal white matter brain adaptations that support fluid intelligence with age

**DOI:** 10.64898/2026.05.27.728295

**Authors:** Josh Neudorf

**Affiliations:** Centre for Social Sciences, Athabasca University, Canada; Department of Psychology, University of Alberta, Canada; Department of Communication Sciences and Disorders, University of Alberta, Canada; Neuroscience and Mental Health Institute, University of Alberta, Canada; Institute for Neuroscience and Neurotechnology, Simon Fraser University, Canada

**Keywords:** . aging brain, diffusion magnetic resonance imaging, network neuroscience, fluid intelligence, physical activity, brain resilience

## Abstract

The aging population continues to grow, and along with it the need to better understand the unique characteristics of the aging brain. Despite the cortical thinning and myelin degeneration that occurs in later life, many individuals are resilient to these changes and are relatively spared from cognitive decline (so called “super-agers”). Longitudinal magnetic resonance imaging data from the Human Connectome Project allowed us to probe the underlying white matter connections that are strengthened or pruned to support fluid intelligence in younger and older adults, while also investigating physical activity as a potential behavioural intervention to promote these network reorganizations. Multivariate partial least squares analysis and the recent method of correlational tractography identified parts of the white matter structural connectivity network, measured with diffusion magnetic resonance imaging, where strengthening or pruning were associated with positive fluid intelligence trajectories and promoted by physical activity. This research improves our understanding of how the brain network adapts in later life, identifies physical activity as an accessible intervention to promote this adaptation, and enriches our fundamental models of what the brain’s neural network is capable of.

---

Considering the rapidly growing older adult population and the unique challenges facing this population, the United Nations has declared the decade from 2021-2030 the Decade of Healthy Aging, calling for action to improve the health and well-being of older adults (United Nations, 2020). One of the challenges facing the aging population is that the aging brain undergoes a host of negative changes, including cortical thinning (Salat et al., 2004) and myelin degradation (Bartzokis, 2004). These brain changes often manifest as cognitive decline, contributing to decreased quality of life. Furthermore, cognitive decline in aging is predictive of future health outcomes (Nelson et al., 2020). However, there are a range of cognitive trajectories across aging individuals, with some being so exceptionally spared that they have been labelled “super-agers” (Harrison et al., 2012, 2018; Mohammadiarvejeh et al., 2022; Zhang et al., 2020). Our recent research has demonstrated that the brain has the remarkable ability to reorganize its structural and functional connectivity networks to compensate for detrimental brain changes (Neudorf et al., 2024, 2025).

The discrepancy between individuals’ cognitive trajectories with age leads to the important question of why some respond poorly while others are spared, and how we can promote brain resilience (see Stern et al., 2023; Udeh-Momoh et al., 2025 for reviews). Among potential behavioural interventions, physical activity (in the form of aerobic exercise) stands out as a particularly attractive recommendation due to its low cost and additional benefits to whole-body health. Animal models have demonstrated that physical activity increases brain growth factors, stimulates neurogenesis, promotes resilience against brain insult, and manifests as better learning and cognitive outcomes (see Cotman & Berchtold, 2002 for a review). Across the lifespan in humans, from childhood to older age, physical activity is associated with better cognitive ability. Furthermore, cortical gray matter volume is larger for individuals with more physical activity, and physical activity modulates brain activity as measured by electroencephalography (EEG) and functional magnetic resonance imaging (fMRI; see Hillman et al., 2008 for a review). Brain network analyses have suggested that physical activity may promote plasticity of the network. For example, in older adults physical activity was associated with increased functional connectivity (from fMRI) in the default mode network and frontal parietal network, and these changes were associated with improved executive function (Voss, Erickson, et al., 2010; Voss, Prakash, et al., 2010). EEG signal analysis using multiscale entropy has demonstrated that physical activity in older adults promotes more local information processing in the brain network and better cognitive ability (Heisz et al., 2015). The effects on the brain’s white matter have also been investigated, as measured with diffusion MRI, demonstrating increase fractional anisotropy in prefrontal and temporal regions (Voss et al., 2013).

The current research aims to investigate the role of the brain’s white matter structural connectivity network in the relationship between age, fluid intelligence, and physical activity. A whole-brain approach utilizing multivariate partial least squares (PLS) will attempt to identify longitudinal structural connectivity network changes associated with physical activity and preserved fluid intelligence, in addition to how this relationship differs as a function of age. A recent method called correlational tractography will also be used to identify specific segments of the white matter tracts whose longitudinal anisotropy changes are associated with physical activity and preserved fluid intelligence. The newly released Human Connectome Project Aging Adult Brain Connectome (HCP – AABC) longitudinal data release will be utilized, containing high quality data with multiple timepoints. Longitudinal data provides valuable insights into individualized aging trajectories, but is often not feasible for practical reasons. Recent research on the aging brain and cognitive trajectories has demonstrated that cross-sectional and longitudinal results may not converge, leading to the recommendation that longitudinal data be utilized whenever possible (Vidal-Pineiro et al., 2021). Furthermore, longitudinal data are important for studying brain resilience, as these data allow us to identify the slopes (changes) of brain and behavioural measures, rather than simply the intercept (initial state) that allows some individuals to grow older before noticing detrimental changes. Our previous research using cross-sectional data did not detect lifestyle factors that supported the aging brain structural connections associated with preserved fluid intelligence (Neudorf et al., 2024), but by using longitudinal data we hypothesize that we should be able to identify adaptive structural connectivity network reorganizations that are promoted by physical activity.

## Methods

Data came from the Human Connectome Project Aging Adult Brain Connectome (HCP – AABC) dataset (https://humanconnectome.org/study/hcp-lifespan-aging), which includes the diffusion MRI data used here to identify white matter connectivity via generalized q-sampling imaging (GQI; Yeh et al., 2010), as well as behavioural measures including fluid intelligence (a fluid IQ factor score based on the NIH Toolbox Dimensional Change Card Sort Test, Flanker Inhibitory Control and Attention Test, Pattern Comparison Test, and Trail Making Test A and B; see Hong et al., 2026; https://nihtoolbox.org/domain/cognition), physical activity (from the International Physical Activity Questionnaire physical activity category summary measure; see Booth, 2000), and age. These data were collected on the same subjects at multiple timepoints, allowing for a longitudinal investigation.

Healthy participants were included based on availability of behavioural and dMRI data. Participants with a diagnosis of mild cognitive impairment or Alzheimer’s Disease were excluded. A total of 715 participants (417 female at birth) were included in the PLS analysis, which included all ages. Of those that responded to the gender identity question, one identified as transgender and one indicated that they did not know what gender they identified with. The mean age was 59.754 years (*SD* = 14.609 years). For the correlational tractography analyses focusing on older adult fluid Intelligence, 497 participants (284 female at birth) were included. One of these participants indicated that they identified as transgender. These participants were older than 50 years old, and the mean age was 67.199 years (SD = 10.796 years). For the correlational tractography analyses focusing on older adult physical activity, 574 participants (328 female at birth) were included. One of these participants indicated that they identified as transgender. These participants were older than 50 years old, and the mean age was 68.021 years (*SD* = 11.048 years). For the correlational tractography analyses focusing on younger adult fluid intelligence, 218 participants (133 female at birth) were included. One of these participants indicated that they did not know which gender they identified with. These participants were younger than or equal to 50 years old, and the mean age was 42.780 years (*SD* = 4.407 years). For the correlational tractography analyses focusing on younger adult physical activity, 241 participants (145 female at birth) were included. One of these participants indicated that they did not know which gender they identified with. These participants were younger than or equal to 50 years old, and the mean age was 42.726 years (*SD* = 4.437 years).

### Diffusion MRI

Preprocessed data from the Human Connectome Project was used in these analyses (see details in the original publication for the first release, Bookheimer et al., 2019). A multishell diffusion scheme was used, and the b-values were 1495 and 2995 s/mm². The number of diffusion sampling directions were 93 and 92, respectively. The in-plane resolution was 1.5 mm. The slice thickness was 1.5 mm. Tractography was performed using DSI Studio (Yeh, 2025), the Hou version (accessed January 30, 2026, http://dsi-studio.labsolver.org). The diffusion data were reconstructed in the MNI space using q-space diffeomorphic reconstruction (Yeh & Tseng, 2011) to obtain the spin distribution function (Yeh et al., 2010). A diffusion sampling length ratio of 1.25 was used. The output resolution in diffeomorphic reconstruction was 1.5 mm isotropic. The restricted diffusion was quantified using restricted diffusion imaging (Yeh et al., 2017). The difference between longitudinal scans were calculated by Visit 2 – Visit 1.

### Partial Least Squares Connectivity Analysis

For the PLS analysis of the structural connectivity matrix, a deterministic fiber tracking algorithm (Yeh et al., 2013) was used with augmented tracking strategies (Yeh, 2020) to improve reproducibility. Using DSI Studio’s default procedure, the anisotropy threshold was randomly selected between 0.5 and 0.7 otsu threshold. The angular threshold was randomly selected from 45 degrees to 90 degrees. The step size was set to voxel spacing. Tracts with length shorter than 30 mm or longer than 200 mm were discarded. A total of 1,000,000 seeds were placed. The connectivity matrix was calculated using a combination of the Schaefer 200 region cortical atlas (Schaefer et al., 2018) and the Tian subcortical atlas (Tian et al., 2020). The subcortical regions included regions from the Tian Scale 1 atlas but with the hippocampus represented by the Tian Scale 3 atlas definition, with the two head divisions combined into a single parcel. These regions were set as “pass” regions in DSI Studio, with whole-brain seeding.

Multivariate partial least squares (PLS) analysis (McIntosh & Lobaugh, 2004) was used to identify latent variables (LVs), each containing weights that describe the relationship of all connections with age, time between visits (days), change in fluid intelligence across visits, and physical activity. PLS uses singular value decomposition to project the data matrix onto orthogonal LVs (similar to principal component analysis). The significance of each identified LV is determined via permutation testing, whereby the order of the data is randomized over a large number of iterations so that the independent and dependent variables are no longer paired to one another, in order to obtain a permutation *p* value representing the significance of that LV. We report only the most reliable PLS weights as determined by bootstrap resampling, which is used to calculate bootstrap ratios (BSR). The bootstrap ratio indicates the reliability of the PLS weights and is calculated as the ratio of the PLS weights (saliences) and their standard errors as determined by bootstrap resampling (Kovacevic et al., 2013; McIntosh & Lobaugh, 2004). The BSRs are analogous to a Z-score, and as such a value exceeding an absolute value of 2 is considered reliable. In these analyses, a conservative threshold value of 4 was used in order to highlight the most reliable results. This procedure was performed using 1,000 iterations for permutation testing and bootstrap resampling. The longitudinal change between the structural connectivity quantitative anisotropy (QA) data in Visit 1 and Visit 2 were given as the independent variables and the dependent variables were Age, longitudinal Days between visits, change in Fluid Intelligence across visits, and Physical Activity.

### Correlational Tractography

Correlational tractography (Yeh et al., 2021) was derived to visualize pathways that have a longitudinal change of QA correlated with the variable of interest (Physical Activity or longitudinal change in Fluid Intelligence) controlling for the time between visits. A nonparametric Spearman partial correlation was used to derive the correlation, and the effect of the number of days between visits was removed using a multiple regression model. The statistical significance of the correlation was examined using a permutation test (Yeh et al., 2016). A t-score threshold of 2 was used in the fiber tracking algorithm (Yeh et al., 2013). A seeding region was placed as the whole brain. The tracks were filtered by topology-informed pruning (Yeh et al., 2019) with 16 iterations. A length threshold of 10 voxel distance was used to select tracks. To estimate the false discovery rate, a total of 4,000 randomized permutations were applied to the group label to obtain the null distribution of the tract length. All reported results had a false discovery rate of less than .001.

## Results

### Partial Least Squares Analysis

The PLS analysis of structural connectivity QA identified two significant LVs. LV1 (permutation *p* < .001) was negatively correlated with Age and longitudinal time between visits (Days) but positively correlated with the change in Fluid Intelligence and the Physical Activity score (see Figure 1A). The positive connections (red) in Figure 1B, then, are those that are associated with better Fluid Intelligence trajectories longitudinally, especially in younger adults, with Physical Activity promoting the strength of these connections and a longer time between visits diminishing the effect. Conversely, the negative connections (blue) indicate those that are reduced (i.e., pruned) to support Fluid Intelligence in younger adults (this pruning promoted by Physical Activity and diminished by time between visits). Of the positive connections, one was within the LH and 16 were interhemispheric. Network regions were involved 10 times from Default Mode, 9 times from Somatomotor, 7 times from Dorsal Attention, 5 times from Control, 1 time from Limbic, 1 time from Salience Ventral Attention, and 1 time from Subcortical. The most highly connected regions are listed in Table 1, and include 3 LH Default Mode PFC regions, a RH PFC region from the Control network, the LH FEF from the Dorsal Attention network, and a LH Somatomotor region. Of the negative connections, 9 were within the LH, 9 were within the RH, and 1 was interhemispheric. Network regions were involved 16 times from Subcortical, 6 times from Visual, 6 times from Somatomotor, 4 times from Control, 3 times from Default Mode, 1 time from Dorsal Attention, 1 time from Salience Ventral Attention, and 1 time from Limbic. The most highly connected regions are listed in Table 1, and include anterior and posterior thalamus regions from both hemispheres, the RH putamen, a RH Visual region, and a LH Somatomotor region.

**Figure 1.**
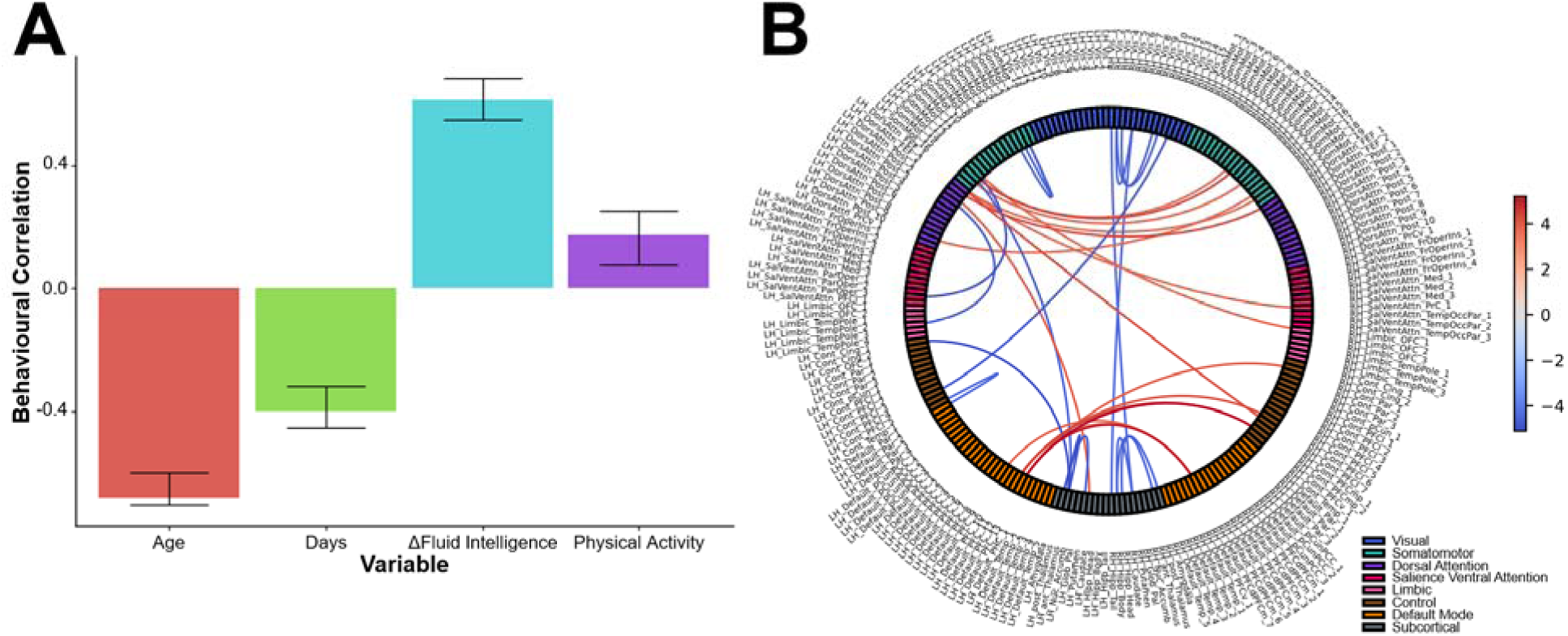
SC PLS analysis latent variable 1 (*p* < .001) behavioural correlation (A) and connection bootstrap ratios (B), indicating the connections reliably correlated negatively with age and longitudinal days, and positively correlated with the longitudinal change in fluid intelligence and physical activity. The colour bar for (B) corresponds to the bootstrap ratio.

**Table 1.** Highest degree regions from the significant network identified by the PLS LV1 of age, longitudinal days, longitudinal change in fluid intelligence, and physical activity (only showing regions with more than 1 connection).

| Region (Positive) | Pos. Degree | Region (Negative) | Neg. Degree |
| --- | --- | --- | --- |
| LH_Default_PFC_13 | 3 | LH_Ant_Thalamus | 3 |
| LH_DorsAttn_FEF_1 | 3 | RH_Post_Thalamus | 3 |
| RH_Default_PFCdPFCm_7 | 3 | LH_Post_Thalamus | 2 |
| LH_Default_PFC_12 | 3 | RH_Ant_Thalamus | 2 |
| LH_SomMot_14 | 2 | RH_Vis_2 | 2 |
| RH_Cont_PFCmp_2 | 2 | LH_SomMot_2 | 2 |
|  |  | RH_Putamen | 2 |

The second LV (permutation *p* = .003) was positively correlated with Age, the change in Fluid Intelligence, and Physical Activity, but negatively correlated with the longitudinal time between visits (Days; see Figure 2A). The positive connections (red) in Figure 2B, then, are those that are associated with better Fluid Intelligence trajectories longitudinally, especially in older adults, with Physical Activity promoting the strength of these connections and a longer time between visits diminishing the effect. Conversely, the negative connections (blue) indicate those that are reduced (i.e., pruned) to support Fluid Intelligence in older adults (this pruning promoted by Physical Activity and diminished by time between visits). Of the positive connections, 5 were within the LH, 2 were within the RH, and 3 were interhemispheric. Network regions were involved 9 times from Subcortical, 5 times from Default Mode, 2 times from Control, 1 time from Visual, 1 time from Somatomotor, 1 time from Dorsal Attention, and 1 time from Limbic. The most highly connected regions were the LH anterior thalamus and the LH hippocampal head, with a degree of 2 for both. Of the negative connections, all 3 were interhemispheric. Network regions were involved 2 times from Somatomotor, 1 time from Dorsal Attention, 1 time from Limbic, 1 time from Control, and 1 time from Default. None of these negative connections had a degree greater than 1.

**Figure 2.**
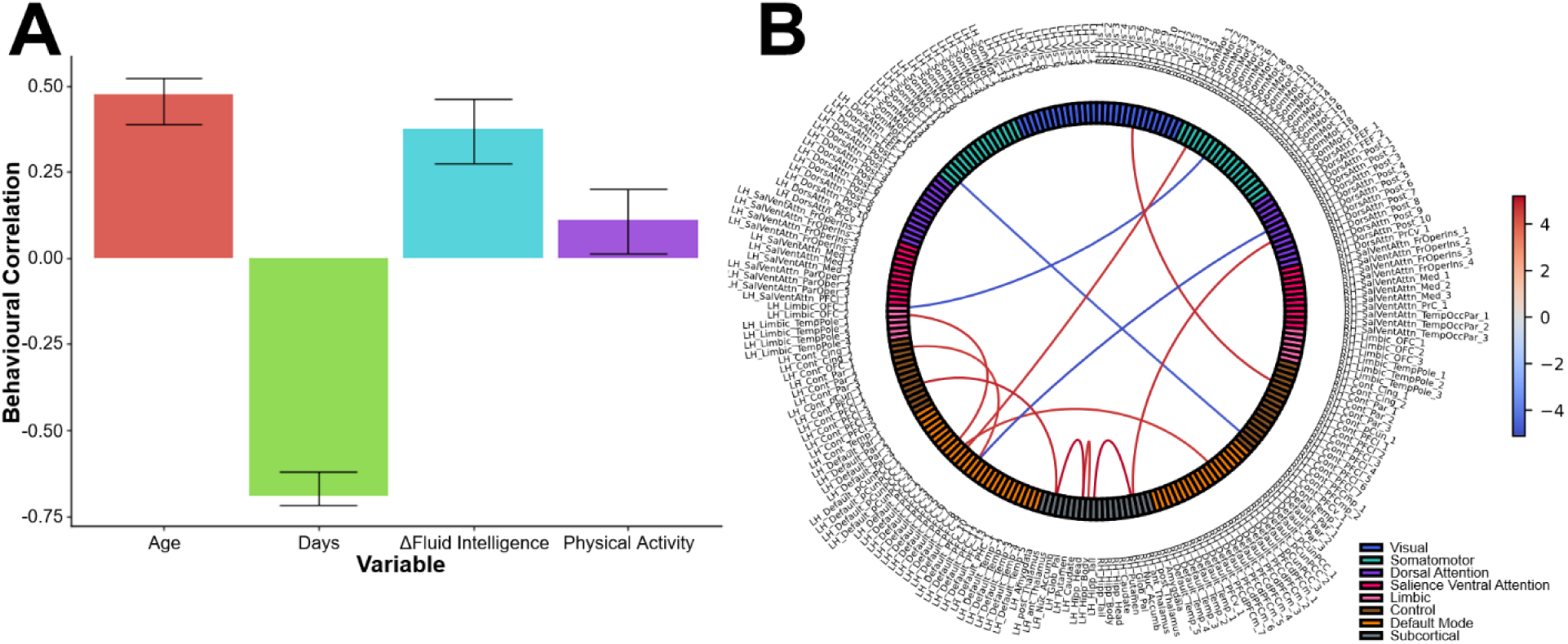
SC PLS analysis latent variable 2 (*p* = .003) behavioural correlation (A) and connection bootstrap ratios (B), indicating the connections reliably correlated negatively with age, longitudinal change in fluid intelligence and physical activity, and positively correlated with longitudinal days. The colour bar for (B) corresponds to the bootstrap ratio.

### Correlational Tractography

The correlational tractography analysis identified longitudinal changes in specific tract segments associated with a longitudinal change in Fluid Intelligence (see Figures 3 and 4). This analysis also identified longitudinal changes in tract segments associated with Physical Activity (see Figures 5 and 6). The number of tracts identified was much higher for older adults than for younger adults in the Fluid Intelligence analysis, and in the Physical Activity analysis an extensive network was identified for older adults while a minimal number of tracts was identified for younger adults. Furthermore, there was extensive overlap in the tracts identified in the Fluid Intelligence and Physical Activity analyses (see Table 2), while younger adults exhibited much less overlap (see Supplementary Table 4). For this reason, the results for the older adults will be focused on in this section, with the younger adult results reported in the Supplementary Materials.

**Figure 3.**
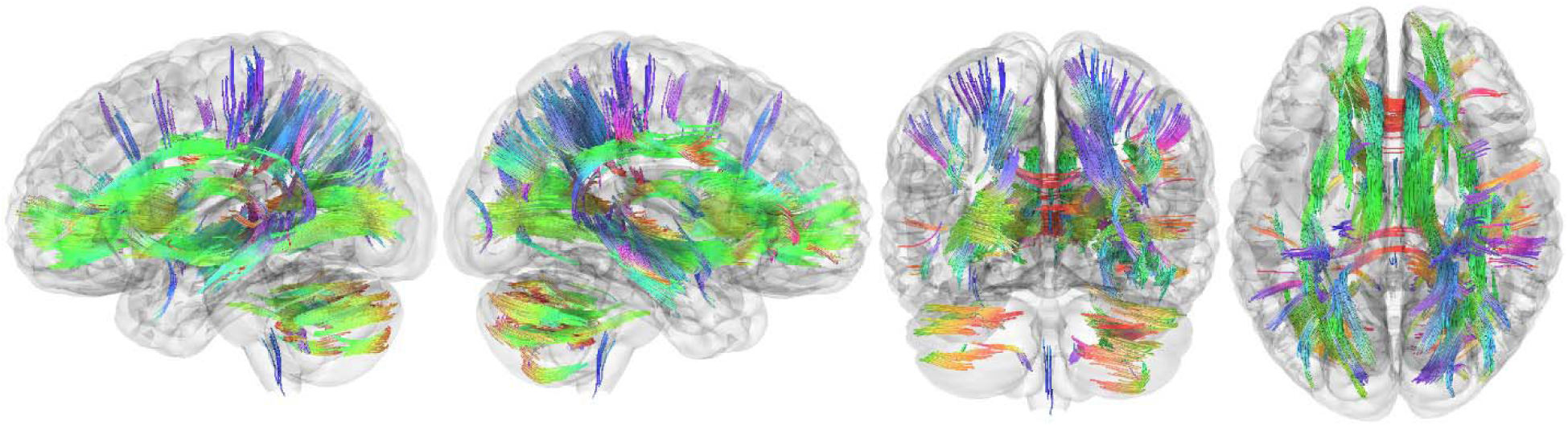
Correlational tractography tracts exhibiting a positive relationship between longitudinal change in structural connectivity and longitudinal change in fluid intelligence, controlling for the number of days between visits, in older adults. Colour indicates tract direction.

**Figure 4.**
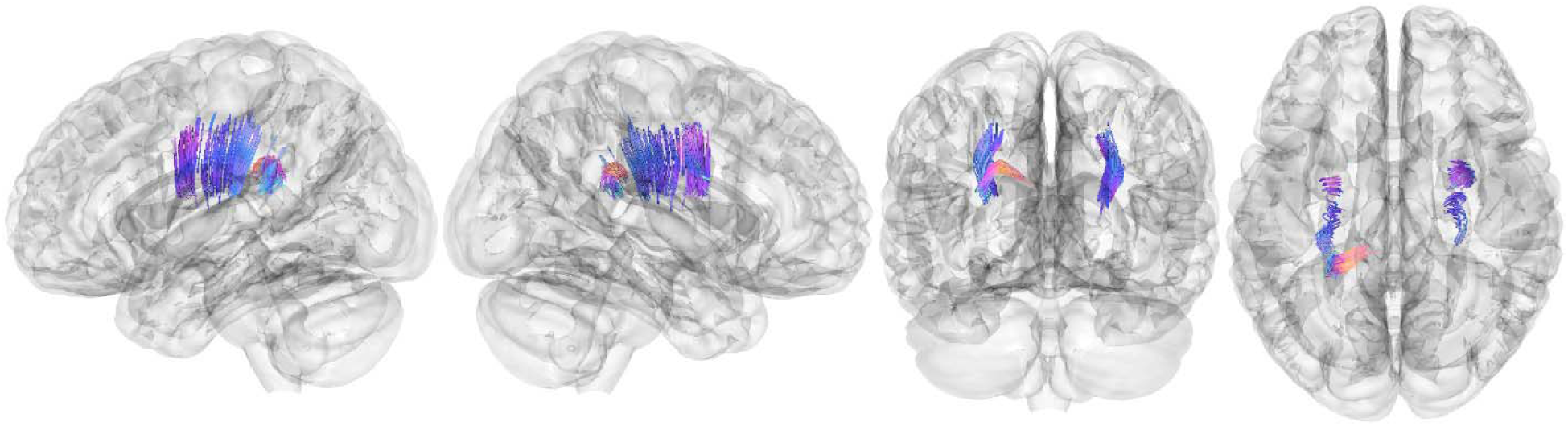
Correlational tractography tracts exhibiting a negative relationship between longitudinal change in structural connectivity and longitudinal change in fluid intelligence, controlling for the number of days between visits, in older adults. Colour indicates tract direction.

**Figure 5.**
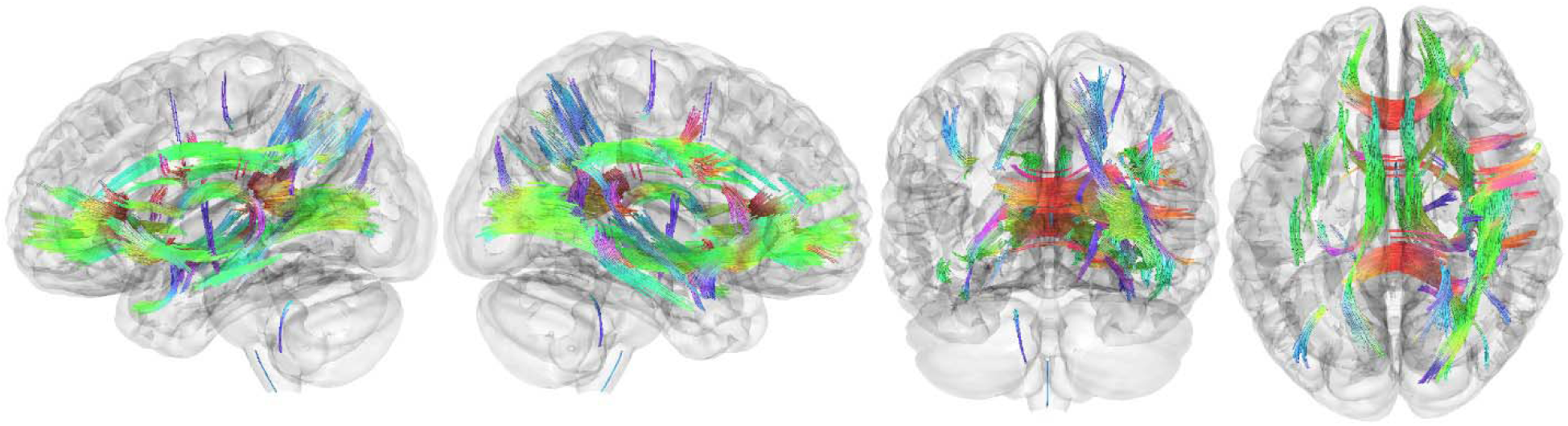
Correlational tractography tracts exhibiting a positive relationship between longitudinal change in structural connectivity and physical activity, controlling for the number of days between visits, in older adults. Colour indicates tract direction.

**Figure 6.**
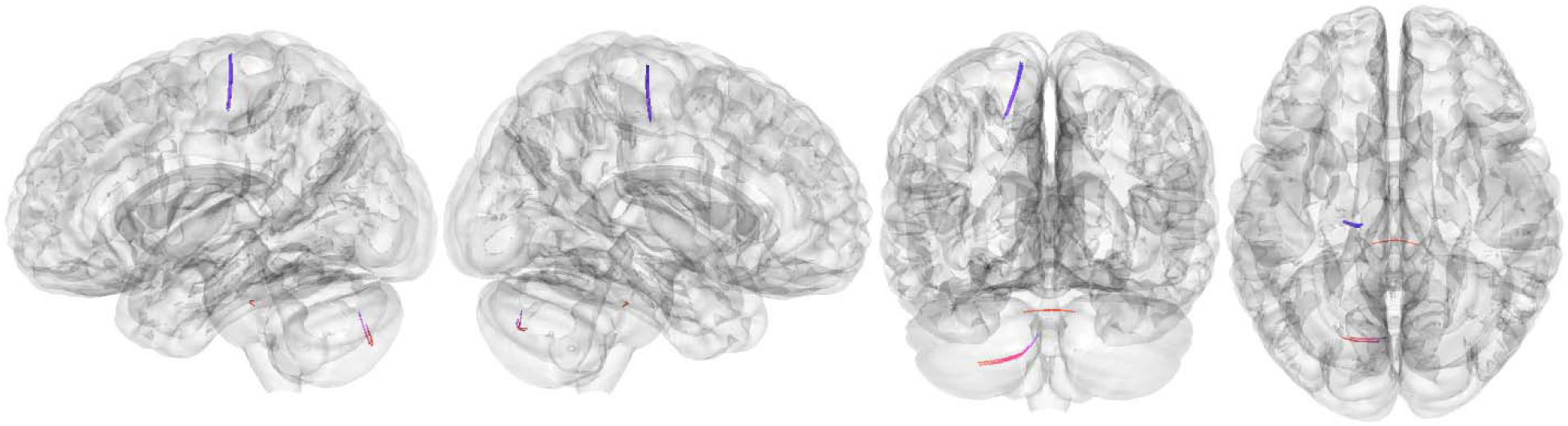
Correlational tractography tracts exhibiting a negative relationship between longitudinal change in structural connectivity and physical activity, controlling for the number of days between visits, in older adults. Colour indicates tract direction.

**Table 2.** Primary tracts identified by correlational tractography as significantly related to both changes in Fluid Intelligence (FI) and Physical Activity (PA) in older adults.

**Positive Correlation**
| Tract Name | FI Tracts | PA Tracts | Total Tracts |
| --- | --- | --- | --- |
| Commissure_CorpusCallosum_ForcepsMajor | 875 | 312 | 1187 |
| Commissure_CorpusCallosum_ForcepsMinor | 415 | 532 | 947 |
| Commissure_CorpusCallosum_Tapetum | 313 | 321 | 634 |
| Association_InferiorLongitudinalFasciculusR | 270 | 235 | 505 |
| ProjectionBasalGanglia_ThalamicRadiationR_Anterior | 398 | 72 | 470 |
| ProjectionBasalGanglia_ThalamicRadiationL_Anterior | 370 | 32 | 402 |
| Association_UncinateFasciculusR | 32 | 292 | 324 |
| Commissure_CorpusCallosum_Body | 243 | 56 | 299 |
| Association_InferiorFrontoOccipitalFasciculusL | 172 | 105 | 277 |
| ProjectionBasalGanglia_ThalamicRadiationR_Superior | 204 | 71 | 275 |

**Negative Correlation**
| Tract Name | FI Tracts | PA Tracts | Total Tracts |
| --- | --- | --- | --- |
| ProjectionBasalGanglia_CorticostriatalTractL_Superior | 3 | 5 | 8 |

For older adults, the most prominent tracts identified as longitudinally related to the change in Fluid Intelligence included the corpus callosum (forceps major and minor, tapetum, and body), RH inferior longitudinal fasciculus, bilateral anterior thalamic radiation, RH superior thalamic radiation, LH inferior fronto-occipital fasciculus, bilateral cerebellum, middle cerebellar peduncle, and cerebellar vermis. With the exception of the cerebellar tracts, all of these prominent tracts were also identified as being stronger in those engaging in more Physical Activity (see Figure 4 and Table 2). In total, there were 52 labelled white matter fibre tracts identified as being associated with both Fluid Intelligence and Physical activity (see Supplementary Table 1). A smaller number of tract segments were identified as decreased in individuals with better Fluid Intelligence trajectories over the same time period, including the bilateral corticospinal tract, RH superior corticostriatal tract, and LH medial lemniscus (see Figure 4 and Supplementary Table 2). Tracts associated only with Physical Activity and not with Fluid Intelligence were very minimal, involving only a handful of fibre tracts (see Supplementary Table 3) indicating that Physical Activity serves to promote the same structural connectivity network changes that support Fluid Intelligence in older age.

## Discussion

These analyses identified many structural connections that change over the lifespan to support fluid intelligence, and that physical activity promotes these changes. The PLS analysis of the parcellated connectivity data identified the most important connection growth and pruning supporting this effect for younger and older adults specifically. For younger adults, the connections that were promoted by physical activity to support better fluid intelligence were mostly interhemispheric, while those that were pruned were mostly intrahemispheric. Conversely, in older adults the connections that were promoted by physical activity to support better fluid intelligence were all intrahemispheric, while those that were pruned were mostly interhemispheric. These findings replicate those we have found previously in a cross-sectional study (Neudorf et al., 2024), and add evidence supporting our theory that older brains rely most on a network reorganization that strengthens local connections to rely less on global connections that are most vulnerable to demyelination (see also Heisz et al., 2015). The default mode network was one of the most highly implicated networks in the connections that were strengthened for both younger and older adults, and of these default mode network connections all involved prefrontal cortical regions. The younger adult pattern of interhemispheric connections between the prefrontal cortex is featured prominently in Figure 1, along with other prominent patterns of interhemispheric connectivity between Somatomotor regions and also between the frontal eye fields of the Dorsal Attention network. Subcortical regions were also highly implicated in this relationship, but these connections were mostly pruned for younger adults and strengthened for older adults. Specifically, younger adults pruned multiple connections to the bilateral anterior and posterior thalamus as well as the RH putamen. Conversely, older adults strengthened multiple connections to the LH anterior thalamus and the LH hippocampal head.

The correlational tractography analyses add specificity to these results, identifying specific longitudinal tract segment changes associated with better fluid intelligence trajectories and physical activity. The tract segments identified showed extensive overlap between the fluid intelligence and physical activity analyses for older adults, with much less overlap for younger adults, suggesting that the benefits of exercise for the identified tracts that support fluid intelligence may be more pronounced in older age. However, the PLS analysis found that physical activity supported fluid intelligence-related connections for both younger and older adults, so it should be emphasized that physical exercise is an important promoting factor across the lifespan, even if it may be particularly important for older adults. The tracts identified as being positively associated with both fluid intelligence trajectory and physical activity for older adults were extensive, and included most prominently the corpus callosum (forceps major and minor, tapetum, and body), RH inferior longitudinal fasciculus, bilateral anterior thalamic radiation, RH superior thalamic radiation, and LH inferior fronto-occipital fasciculus. A smaller number of tract segments including the LH superior corticostriatal tract were negatively associated with fluid intelligence trajectory and physical activity, suggesting that pruning in these tracts benefit fluid intelligence and this pruning is promoted by physical activity.

## Conclusion

This research adds further evidence to the theory that aging brain changes are not all negative. In healthy adults, aging brain connectivity adapts by strengthening and pruning connections to reconfigure the network in a way that preserves fluid intelligence. Demonstrating this effect longitudinally strengthens the cross-sectional evidence we have contributed in favour of this theory (Neudorf et al., 2024, 2025) and answers recent calls to utilize longitudinal data to investigate brain changes with age whenever possible (Vidal-Pineiro et al., 2021). Furthermore, these findings demonstrate that physical activity promotes the same brain network reorganization that supports fluid intelligence, representing an important behavioural intervention that can improve brain resilience for the aging population. This research answers the United Nations Decade of Healthy Aging call (United Nations, 2020) promoting research that can produce recommendations to improve health and well-being in the growing older adult population.

## Acknowledgements

Data were provided by the Human Connectome Project, WU-Minn Consortium (Principal Investigators: David Van Essen and Kamil Ugurbil; 1U54MH091657) funded by the 16 NIH Institutes and Centers that support the NIH Blueprint for Neuroscience Research; and by the McDonnell Center for Systems Neuroscience at Washington University. This research was enabled in part by support provided by the British Columbia DRI Group and the Digital Research Alliance of Canada (alliancecan.ca), in addition to computational resources generously shared by Prof. Ron Borowsky (NSERC DG 18968-2013-26). I thank research assistant Melissa Fontaine for her review of the literature, which supported this research.

## Data Availability Statement

Data are available from the Human Connectome Project Aging Adult Brain Connectome (HCP – AABC) dataset (https://humanconnectome.org/study/hcp-lifespan-aging).

Code used to produce these analyses are available in the following repository: https://github.com/neudorf/HCP-AABC_longitudinal_SC

## Supplementary Materials

**Supplementary Table 1.** Structural connectivity tracts identified by correlational tractography as significantly related to both changes in Fluid Intelligence (FI) and Physical Activity (PA) in older adults.

**Positive Correlation**
| Tract Name | FI<br>Tracts | PA<br>Tracts | Total<br>Tracts |
| --- | --- | --- | --- |
| Commissure_CorpusCallosum_ForcepsMajor | 875 | 312 | 1187 |
| Commissure_CorpusCallosum_ForcepsMinor | 415 | 532 | 947 |
| Commissure_CorpusCallosum_Tapetum | 313 | 321 | 634 |
| Association_InferiorLongitudinalFasciculusR | 270 | 235 | 505 |
| ProjectionBasalGanglia_ThalamicRadiationR_Anterior | 398 | 72 | 470 |
| ProjectionBasalGanglia_ThalamicRadiationL_Anterior | 370 | 32 | 402 |
| Association_UncinateFasciculusR | 32 | 292 | 324 |
| Commissure_CorpusCallosum_Body | 243 | 56 | 299 |
| Association_InferiorFrontoOccipitalFasciculusL | 172 | 105 | 277 |
| ProjectionBasalGanglia_ThalamicRadiationR_Superior | 204 | 71 | 275 |
| Association_HippocampusAlveusR | 178 | 96 | 274 |
| Association_CingulumR_FrontalParietal | 175 | 71 | 246 |
| Association_SuperiorLongitudinalFasciculusR_3 | 90 | 147 | 237 |
| Association_CingulumL_FrontalParietal | 129 | 81 | 210 |
| ProjectionBasalGanglia_ThalamicRadiationR_Posterior | 129 | 70 | 199 |
| ProjectionBasalGanglia_FornixL | 101 | 74 | 175 |
| ProjectionBasalGanglia_FornixR | 102 | 73 | 175 |
| Association_CingulumL_Parolfactory | 118 | 44 | 162 |
| Association_SuperiorLongitudinalFasciculusR_2 | 83 | 65 | 148 |
| Association_CingulumR_ParahippocampalParietal | 113 | 33 | 146 |
| Commissure_AnteriorCommissure_Occipital | 49 | 95 | 144 |
| ProjectionBasalGanglia_CorticostriatalTractR_Posterior | 109 | 33 | 142 |
| Association_HippocampusAlveusL | 112 | 21 | 133 |
| ProjectionBasalGanglia_ThalamicRadiationL_Superior | 109 | 14 | 123 |
| ProjectionBasalGanglia_CorticostriatalTractL_Anterior | 104 | 7 | 111 |
| ProjectionBasalGanglia_OpticRadiationL | 96 | 5 | 101 |
| ProjectionBasalGanglia_OpticRadiationR | 39 | 57 | 96 |
| ProjectionBasalGanglia_CorticostriatalTractR_Anterior | 42 | 50 | 92 |
| Association_CingulumR_SuperiorLongitudinalFasciculus1 | 85 | 1 | 86 |
| Association_ArcuateFasciculusR | 39 | 42 | 81 |
| Commissure_AnteriorCommissure_Frontal | 36 | 28 | 64 |
| Association_UncinateFasciculusL | 15 | 46 | 61 |
| Association_ParietalAslantTractL | 39 | 19 | 58 |
| ProjectionBasalGanglia_CorticostriatalTractL_Posterior | 48 | 8 | 56 |
| Association_InferiorLongitudinalFasciculusL | 30 | 24 | 54 |
| ProjectionBrainstem_ReticularTractL | 38 | 12 | 50 |
| Commissure_AnteriorCommissure_Temporal | 2 | 47 | 49 |
| Association_CingulumL_ParahippocampalParietal | 20 | 23 | 43 |
| ProjectionBrainstem_CorticospinalTractR | 15 | 28 | 43 |
| Association_MiddleLongitudinalFasciculusR | 18 | 24 | 42 |
| Association_CingulumL_FrontalParahippocampal | 19 | 18 | 37 |
| ProjectionBrainstem_ReticularTractR | 29 | 7 | 36 |
| Association_FrontalAslantTractR | 8 | 17 | 25 |
| Association_ArcuateFasciculusL | 20 | 2 | 22 |
| ProjectionBasalGanglia_ThalamicRadiationL_Posterior | 16 | 5 | 21 |
| Association_CingulumL_SuperiorLongitudinalFasciculus1 | 4 | 15 | 19 |
| ProjectionBasalGanglia_CorticostriatalTractR_Superior | 12 | 3 | 15 |
| ProjectionBrainstem_CorticobulbarTractR | 9 | 5 | 14 |
| Cerebellum_InferiorCerebellarPeduncleL | 8 | 4 | 12 |
| ProjectionBrainstem_MedialLemniscusR | 9 | 2 | 11 |
| ProjectionBrainstem_CorticopontineTractR_Frontal | 2 | 4 | 6 |
| ProjectionBrainstem_CorticopontineTractR_Occipital | 1 | 2 | 3 |
| **Negative Correlation** |  |  |  |
| Tract Name | FI<br>Tracts | PA<br>Tracts | Total<br>Tracts |
| ProjectionBasalGanglia_CorticostriatalTractL_Superior | 3 | 5 | 8 |

**Supplementary Table 2.** Structural connectivity tracts identified by correlational tractography as significantly related to only changes in Fluid Intelligence (FI) in older adults.

**Positive Correlation**
|  |  |
| --- | --- |
| Tract Name | FI Tracts |
| Cerebellum_CerebellumR | 524 |
| Cerebellum_MiddleCerebellarPeduncle | 373 |
| Cerebellum_CerebellumL | 322 |
| Cerebellum_Vermis | 119 |
| Association_VerticalOccipitalFasciculusR | 24 |
| Cerebellum_InferiorCerebellarPeduncleR | 17 |
| ProjectionBrainstem_NonDecussatingDentatorubrothalamicTractR | 14 |
| Association_ExtremeCapsuleL | 12 |
| Association_SuperiorLongitudinalFasciculusL_2 | 10 |
| ProjectionBrainstem_CorticospinalTractL | 9 |
| ProjectionBrainstem_CorticopontineTractL_Occipital | 8 |
| Association_MiddleLongitudinalFasciculusL | 7 |
| ProjectionBasalGanglia_CorticostriatalTractL_Superior | 7 |
| ProjectionBrainstem_CorticopontineTractR_Parietal | 5 |
| Association_CingulumR_Parahippocampal | 2 |
| Association_SuperiorLongitudinalFasciculusL_3 | 2 |
| ProjectionBrainstem_CorticopontineTractL_Frontal | 2 |
| Cerebellum_SuperiorCerebellarPeduncle | 1 |
| **Negative Correlation** |  |
| Tract Name | FI Tracts |
| Commissure_CorpusCallosum_Tapetum | 78 |
| ProjectionBrainstem_CorticospinalTractR | 69 |
| ProjectionBasalGanglia_CorticostriatalTractR_Superior | 50 |
| ProjectionBrainstem_MedialLemniscusL | 23 |
| ProjectionBrainstem_CorticospinalTractL | 19 |
| ProjectionBrainstem_CorticopontineTractL_Parietal | 9 |
| ProjectionBrainstem_MedialLemniscusR | 6 |
| Association_ExtremeCapsuleL | 5 |
| ProjectionBrainstem_ReticularTractR | 5 |
| ProjectionBasalGanglia_ThalamicRadiationL_Posterior | 3 |
| ProjectionBrainstem_CorticopontineTractR_Parietal | 3 |
| ProjectionBrainstem_NonDecussatingDentatorubrothalamicTractR | 3 |
| ProjectionBasalGanglia_ThalamicRadiationR_Superior | 2 |
| ProjectionBrainstem_CorticopontineTractR_Frontal | 2 |
| Association_FrontalAslantTractR | 1 |
| Commissure_CorpusCallosum_Body | 1 |

**Supplementary Table 3.**
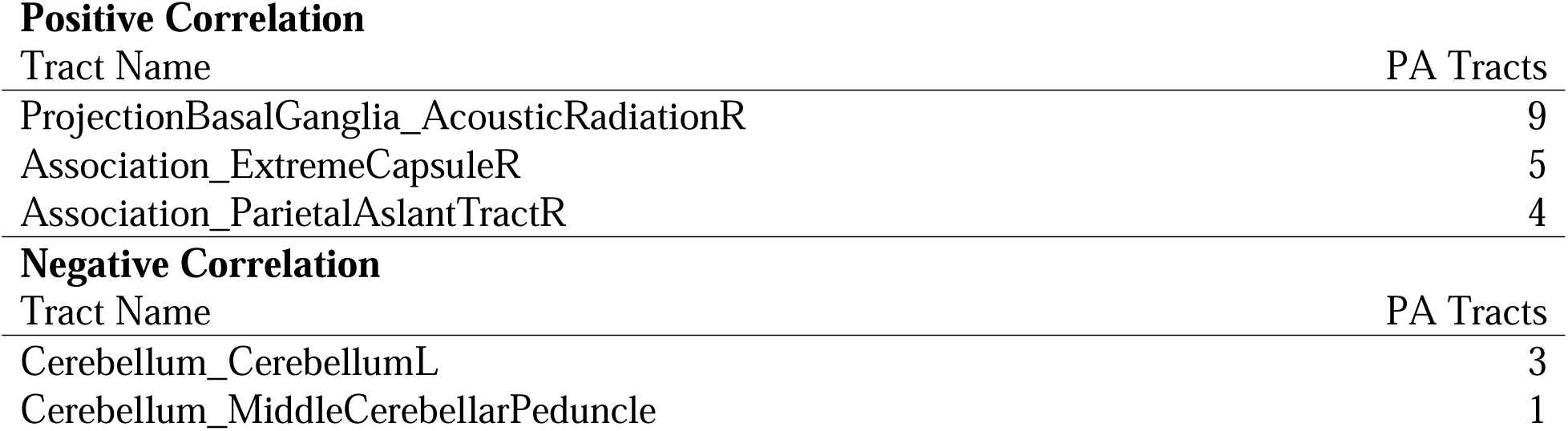
Structural connectivity tracts identified by correlational tractography as significantly related to only Physical Activity (PA) in older adults.

**Supplementary Figure 1.**
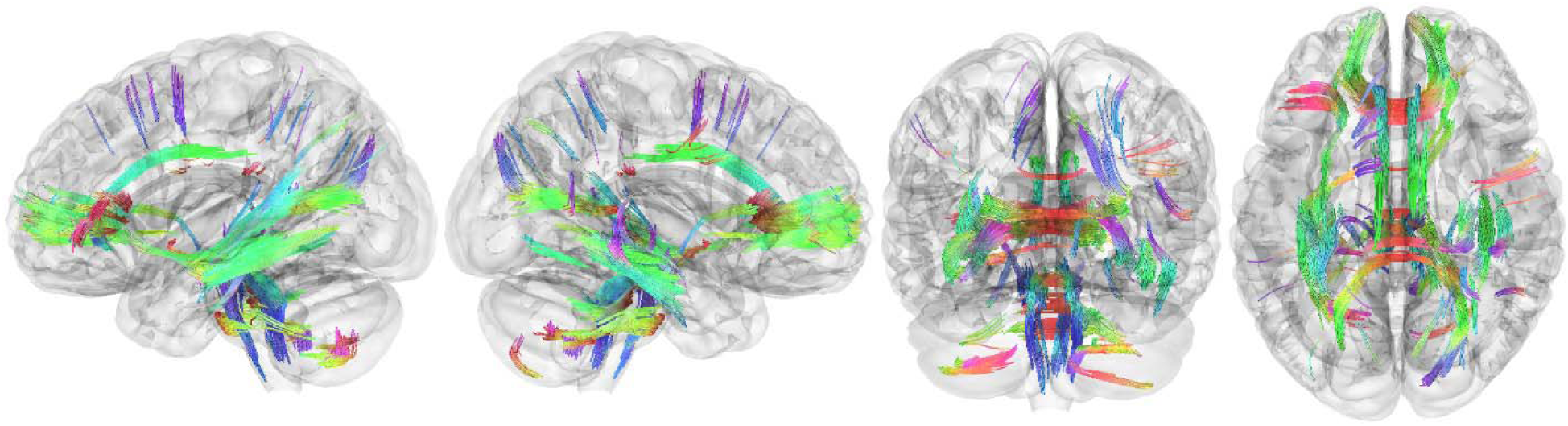
Correlational tractography tracts exhibiting a positive relationship between longitudinal change in structural connectivity and longitudinal change in Fluid Intelligence, controlling for the number of days between visits, in younger adults.

**Supplementary Figure 2.**
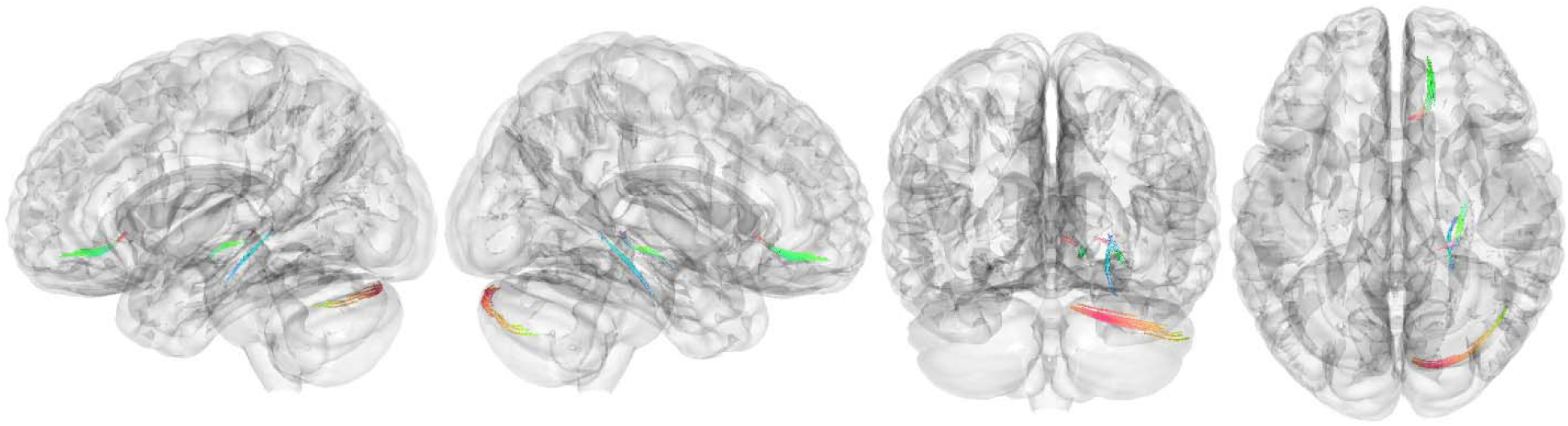
Correlational tractography tracts exhibiting a positive relationship between longitudinal change in structural connectivity and Physical Activity, controlling for the number of days between visits, in younger adults.

**Supplementary Figure 3.**
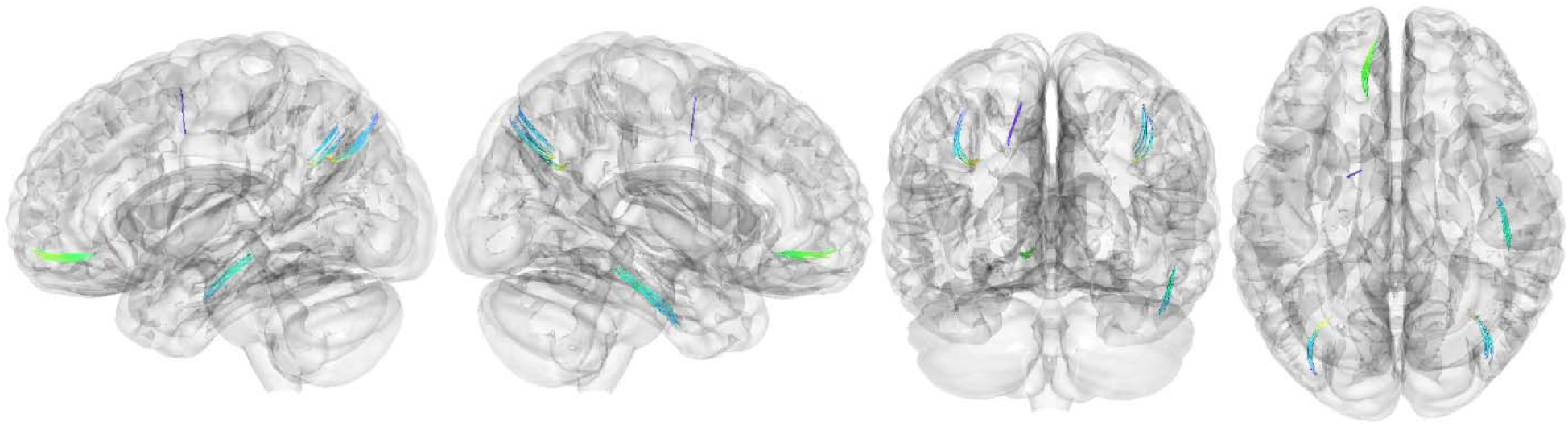
Correlational tractography tracts exhibiting a negative relationship between longitudinal change in structural connectivity and Physical Activity, controlling for the number of days between visits, in younger adults.

**Supplementary Table 4.** Structural connectivity tracts identified by correlational tractography as significantly related to both changes in Fluid Intelligence (FI) and Physical Activity (PA) in younger adults.

**Positive Correlation**
| Tract Name | FI<br>Tracts | PA<br>Tracts | Total<br>Tracts |
| --- | --- | --- | --- |
| Commissure_CorpusCallosum_ForcepsMinor | 534 | 15 | 549 |
| Association_HippocampusAlveusR | 92 | 11 | 103 |
| Association_CingulumR_ParahippocampalParietal | 30 | 13 | 43 |
| ProjectionBasalGanglia_CorticostriatalTractR_Anterior | 33 | 5 | 38 |
| Cerebellum_CerebellumR | 12 | 22 | 34 |
| ProjectionBasalGanglia_FornixR | 22 | 10 | 32 |
| Association_UncinateFasciculusR | 20 | 3 | 23 |

**Supplementary Table 5.**
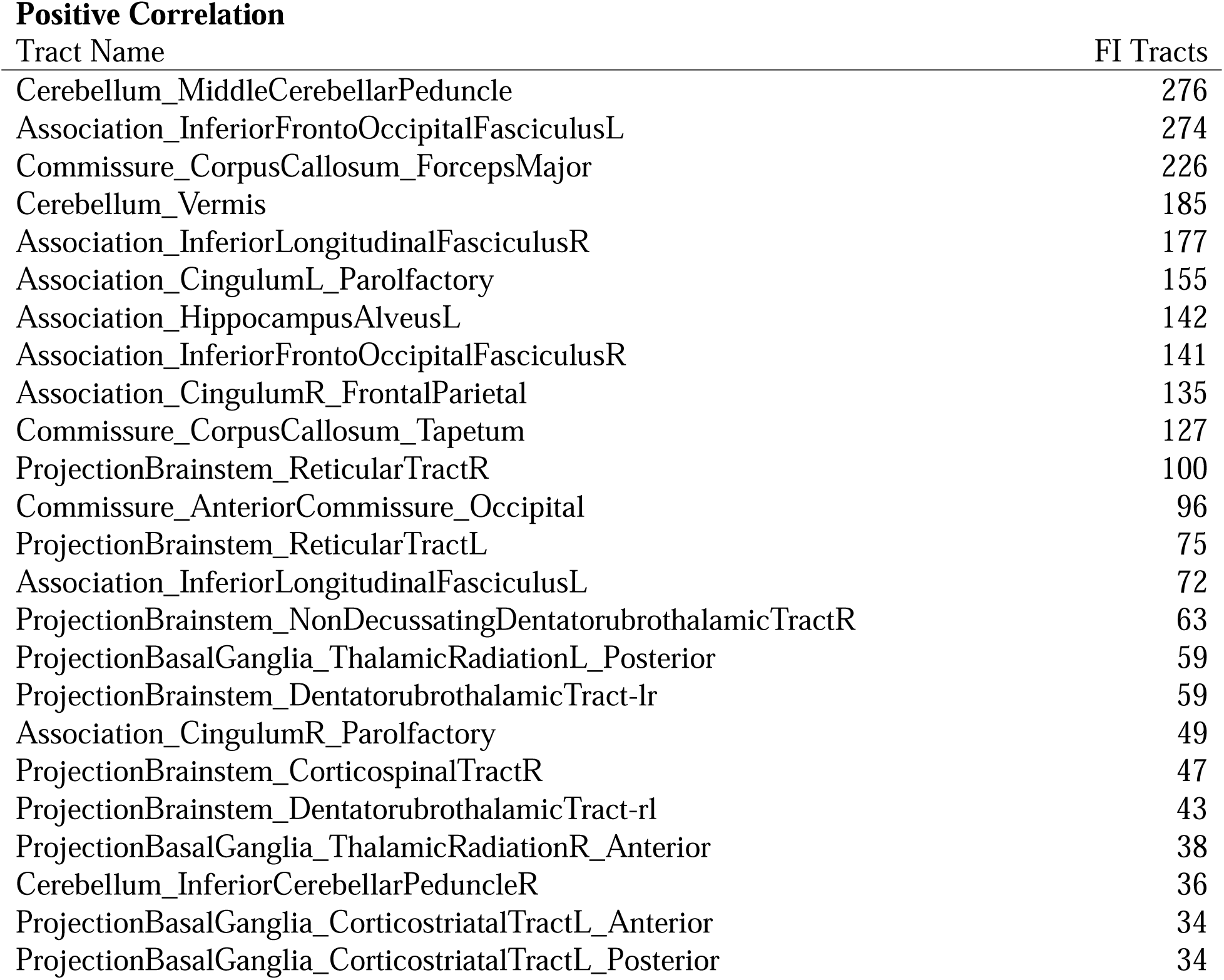
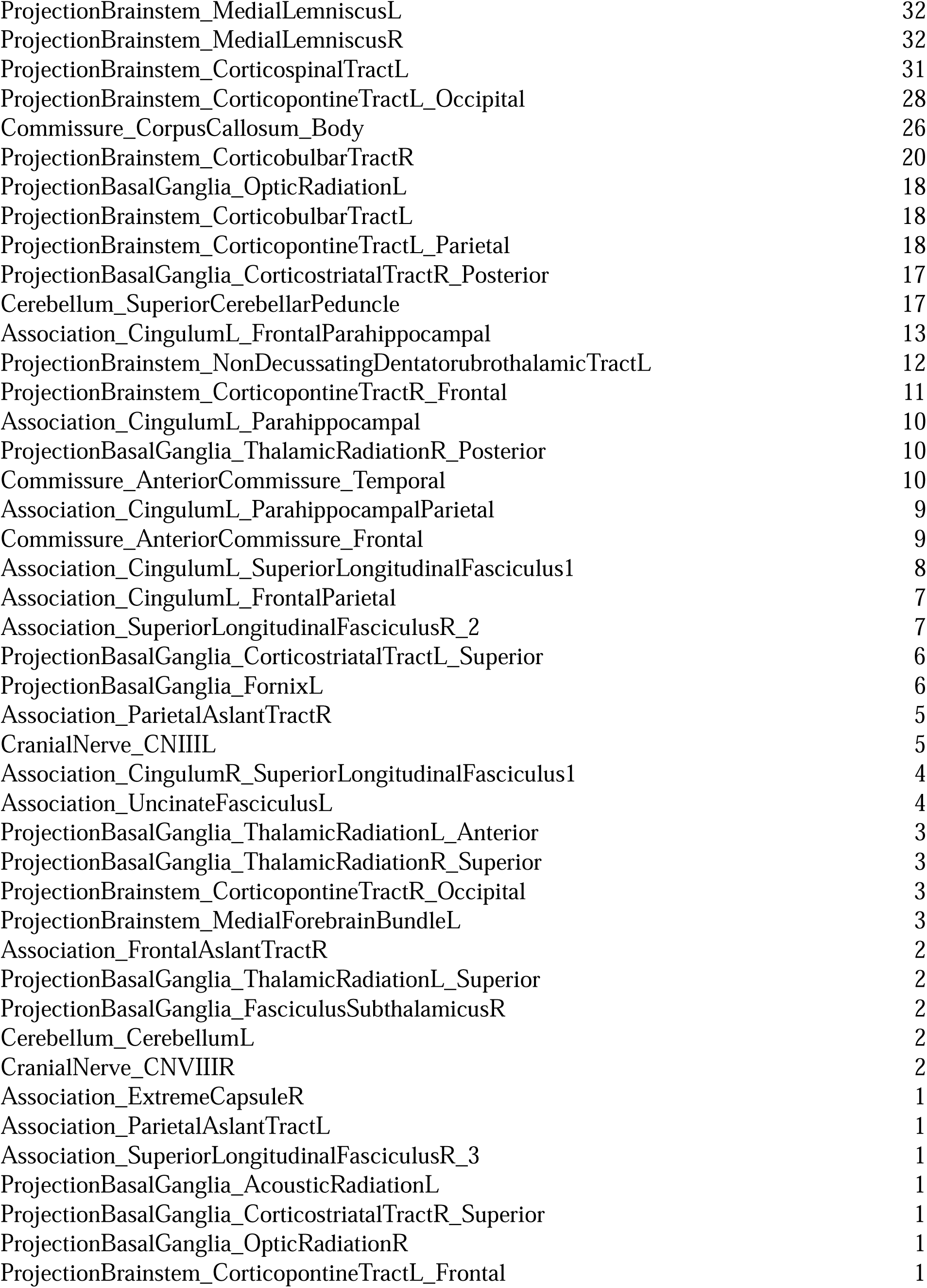
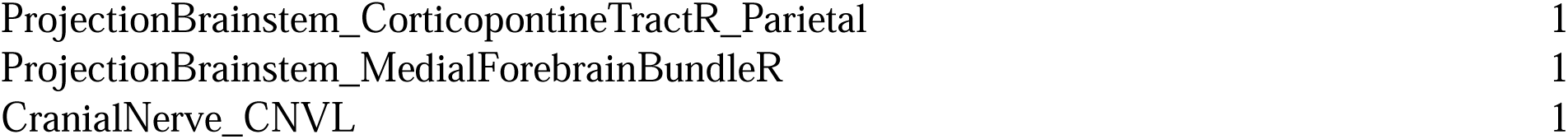
Structural connectivity tracts identified by correlational tractography as significantly related to only changes in Fluid Intelligence (FI) in younger adults.

**Supplementary Table 6.** Structural connectivity tracts identified by correlational tractography as significantly related to only Physical Activity (PA) in younger adults.

**Positive Correlation**
| Tract Name | PA Tracts |
| --- | --- |
| CranialNerve_CNIIR | 2 |

**Negative Correlation**
| Tract Name | PA Tracts |
| --- | --- |
| Commissure_CorpusCallosum_ForcepsMinor | 9 |
| Commissure_CorpusCallosum_Tapetum | 8 |
| ProjectionBasalGanglia_CorticostriatalTractR_Posterior | 7 |
| Association_InferiorLongitudinalFasciculusR | 6 |
| Association_ParietalAslantTractL | 4 |
| ProjectionBasalGanglia_CorticostriatalTractL_Posterior | 3 |
| ProjectionBasalGanglia_ThalamicRadiationL_Anterior | 3 |
| Commissure_CorpusCallosum_ForcepsMajor | 3 |
| ProjectionBasalGanglia_ThalamicRadiationL_Posterior | 1 |
| ProjectionBasalGanglia_ThalamicRadiationR_Posterior | 1 |
| ProjectionBrainstem_CorticopontineTractL_Frontal | 1 |
| Commissure_CorpusCallosum_Body | 1 |

